# Robust identification of extrachromosomal DNA and genetic variants using multiple genetic abnormality sequencing (MGA-Seq)

**DOI:** 10.1101/2022.11.18.517160

**Authors:** Da Lin, Yanyan Zou, Jinyue Wang, Qin Xiao, Fei Lin, Ningyuan Zhang, Zhaowei Teng, Shiyi Li, Yongchang Wei, Fuling Zhou, Rong Yin, Siheng Zhang, Chengchao Wu, Jing Zhang, Sheng Hu, Shuang Dong, Xiaoyu Li, Shengwei Ye, Haixiang Sun, Gang Cao

**Author notes:** These authors contributed equally: Da Lin, Yanyan Zou. Correspondence: Da Lin, Ph.D, Haixiang Sun, Ph.D, Professor, Gang Cao, Ph.D, Professor.

## Abstract

Genomic abnormalities, including structural variation (SV), copy number variation (CNV), single-nucleotide polymorphism (SNP), homogenously staining regions (HSR) and extrachromosomal DNA (ecDNA), are strongly associated with cancer, rare diseases and infertility. A robust technology to simultaneously detect these genomic abnormalities is highly desired for clinical diagnosis and basic research. In this study, we developed a simple and cost-effective method – multiple genetic abnormality sequencing (MGA-Seq) – to simultaneously detect SNPs, CNVs, SVs, ecDNA and HSRs in a single tube. This method has been successfully applied in both cancer cell lines and clinical tumour samples and revealed that focal amplification in tumour tissue is substantially heterogeneous. Notably, we delineated the architecture of focal amplification and the ecDNA network by MGA-Seq, which facilitated the exploration of the regulation of gene expression in ecDNA. This method could be extensively applied for diagnosis and may greatly facilitate the investigation of the genomic mechanism for genetic diseases.

## INTRODUCTION

Genomic abnormalities, including structural variation (SV), copy number variation (CNV), focal amplification (FA) (*1*), and single-nucleotide polymorphisms (SNPs), are strongly associated with the development and progression of cancer (*2, 3*), rare diseases (RDs) (*4*) and infertility (*5, 6*). Accumulating data have demonstrated that numerous cancer cells contain extrachromosomal DNA (ecDNA), a form of FA (*7*). The copy number of oncogenes can be highly elevated by ecDNA-based amplification. Moreover, the chromatin architecture of ecDNA is usually highly accessible (*8*), which dramatically increases the expression level of oncogenes. ecDNAs can be spatially close to each other and cluster together to form ecDNA hubs (*8–10*), which perform enhancer-like functions and increase the expression of proto-oncogenes through intermolecular interactions (*8, 9, 11*). Intriguingly, in response to antitumor drug treatment, ecDNA can reintegrate back into the chromosome in another form of FA, homogenously staining regions (HSRs), via a myriad of mechanisms (*12*). Increasing evidence suggests that ecDNA is associated with cancer progression and can be used as a diagnostic marker (*13, 14*). However, there is no method thus far to simultaneously detect diverse types of genomic abnormalities, which greatly hampers the precise diagnosis and understanding of the molecular mechanism of cancer and genetic disease.

Second-generation sequencing-based whole exome sequencing (WES) and whole genome sequencing (WGS) can efficiently detect single-nucleotide variants (SNVs) and small indels (< 50 bp). However, due to the limitations of short read length, it is extremely challenging to identify larger inversions, translocations, insertions (>1 Mb) (*15*), and ecDNA. To improve the detection capability of complex genomic structural variation, several new technologies have been developed (*15, 16*). These technologies can be generally divided into two categories: one is based on single molecule long fragment sequencing or detection, such as Pacific Biosciences (PacBio) SMRT sequencing (*17, 18*), Oxford Nanopore Technologies (ONT) sequencing (*16, 19, 20*), and Bionano (*21*); the other is based on long DNA sequence reconstruction using short read sequencing, such as strand-seq (*22, 23*), 10x Genomics linked-reads (*24–26*), and Hi-C (*27, 28*). Due to the high cost, tedious experimental steps, and large amount of initial sample, these technologies are mostly applied in scientific research, such as genome assembly (*29–31*), full-length transcriptome sequencing (*32*), and gene transcription regulation (*33*), but not for clinic diagnosis.

Based on WGS datasets, researchers developed the FA prediction software AmpliconArchitect (*34*) and delineated the focal amplifications and general structure of ecDNA in different types of tumours (*35*). Due to the natural disadvantage of the short read length of next-generation sequencing datasets, the accuracy of AmpliconArchitect prediction results is limited, and there is no spatial structural information of ecDNA hubs. Recently, a multiomics strategy based on second-generation sequencing, third-generation sequencing, and Hi-C has been developed to decode the spatial architecture of ecDNA hubs in detail (*9, 36*). This integrated analysis strategy can effectively decode the circular structure and spatial mobility of ecDNA. However, this strategy requires expensive multiple sequencing library construction and sequencing from the same sample, which limits its clinical application for precise diagnosis. Thus, a simple method for the simultaneous detection of different types of genomic abnormalities is crucial and highly desired for precise diagnosis and understanding the molecular mechanism of cancer and genetic disease.

In this study, we sought to develop an efficient and cost-effective method, multiple genetic abnormality sequencing (MGA-Seq), to simultaneously detect SNPs, CNVs, SVs, and the spatial architecture of FA and distinguish ecDNA from HSR. Using MGA-Seq, we successfully identified SNPs, CNVs, specific chromosomal translocation types and breakpoints with single-base resolution in cancer cells and blood samples from infertile patients. As MGA-Seq can locate the approximate location of genomic structural variation, it can facilitate breakpoint searching. We demonstrated that MGA-Seq can indeed distinguish HSR and ecDNA and construct spatial structure and interaction networks of focal amplification regions that could be extensively applied for precise diagnosis and the investigation of the molecular mechanism of cancer and genetic disease.

## Results

### Overview of MGA-Seq

To maintain the spatial architecture of the genome, the nuclei are first fixed by formaldehyde in multiple genetic abnormality sequencing (MGA-Seq). The genome is digested *in situ* by restriction endonuclease followed by 5’ DNA overhang fill-in by DNA polymerase I. Next, the spatially adjacent chromatin fragments are proximity ligated using T4 DNA ligase and then fragmented into a high-throughput sequencing library (**Fig. 1A**). This library contains two kinds of sequencing reads. The reads without proximity ligation junctions were used to detect SNPs, CNVs, small inserts and deletions (< 50 bp), focal amplification (FA), and genomic breakpoints (**Fig. 1A** and **fig. S1**). As the reads with proximity ligation junctions contain spatially adjacent chromosome fragment contact information of the genome, they can be used to decode chromosome structure. Thus, the integrated analysis of all the sequencing reads can identify large chromosome structural variation, such as balanced and unbalanced translocations, extrachromosomal DNA (ecDNA), and intrachromosomal homogenously staining regions (HSRs). Notably, all MGA-Seq steps are carried out in the same tube and do not require buffer replacement, which takes only 9 hours and costs just 56 dollars (**Fig. 1B**).

**Figure 1.**
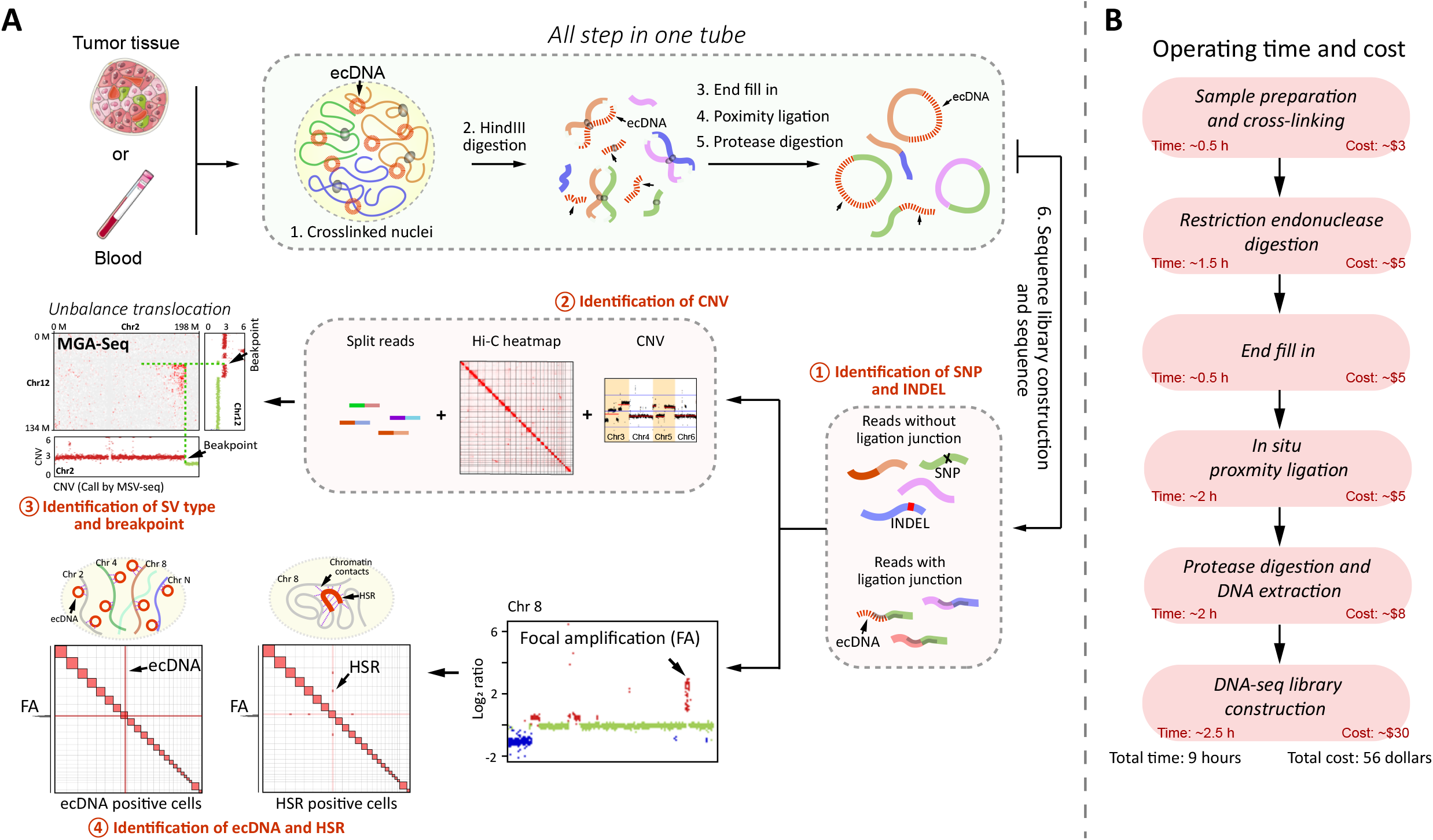
Experimental procedure, time, and cost of multiple genetic abnormalities sequencing (MGA-Seq). **(A)** Flowchart of MGA-Seq. Nuclei were cross-linked with 0.5% formaldehyde and then digested with HindIII. 5’ DNA overhangs of digested chromatin fragments were filled in by DNA polymerase and then proximity ligated by T4 DNA ligase. The proximity ligation products were fragmented and then subjected to high-throughput sequencing library construction. After sequencing, all the reads were used to generate chromatin contact matrix for genome structural variation calling. In the sequencing library, the reads without ligation junction “AAGCTAGCTT” were used for the detection of CNV, SNP, small indels (< 50bp), region of focal amplification, and genome breakpoints. By combining all information, the types and breakpoints of structural variation can be decoded. Notably, MGA-Seq can distinguish ecDNA and HSR, predict the structure of simple focal amplification regions, and construct the interaction network of focal amplificated genes. **(B)** The main steps, time, and cost of MGA-Seq.

### Identification of SNPs and indels by MGA-Seq

To evaluate the SNP and indel detection capability, we performed MGA-Seq on the colorectal cancer cell line SW480 as described in **Fig. 1A**. After sequencing, we obtained 194,167,430 read pairs, of which 2,982,113 (1.5%) read pairs contained “AAGCTAGCTT” ligation junction sequences. To avoid false-positives caused by ligation junctions, we filtered out this part of the reads for SNP and indel detection (see Methods) and analysed the remaining reads by the Genome Analysis Toolkit (GATK). To evaluate the SNP and indel variation calling efficacy, we used the SW480 cell line to generate standard WGS datasets and downloaded the SW480 *in situ* Hi-C datasets (*37*) for comparison with the same parameters (see Methods). As shown in **Fig. 2A**, MGA-Seq identified 2,722,682 variants, including 2,446,823 SNPs, 130,087 insertions, and 145,772 deletions. A total of 82.8% of these variants were consistent with WGS (**Fig. 2B**). Hi-C found only 1,166,315 variants, which is much lower than that identified by MGA-Seq and WGS (**Fig. 2A**). Furthermore, the sequencing coverage and depth of MGA-Seq were also much higher than those of Hi-C (**fig. S2, A-C**).

**Figure 2.**
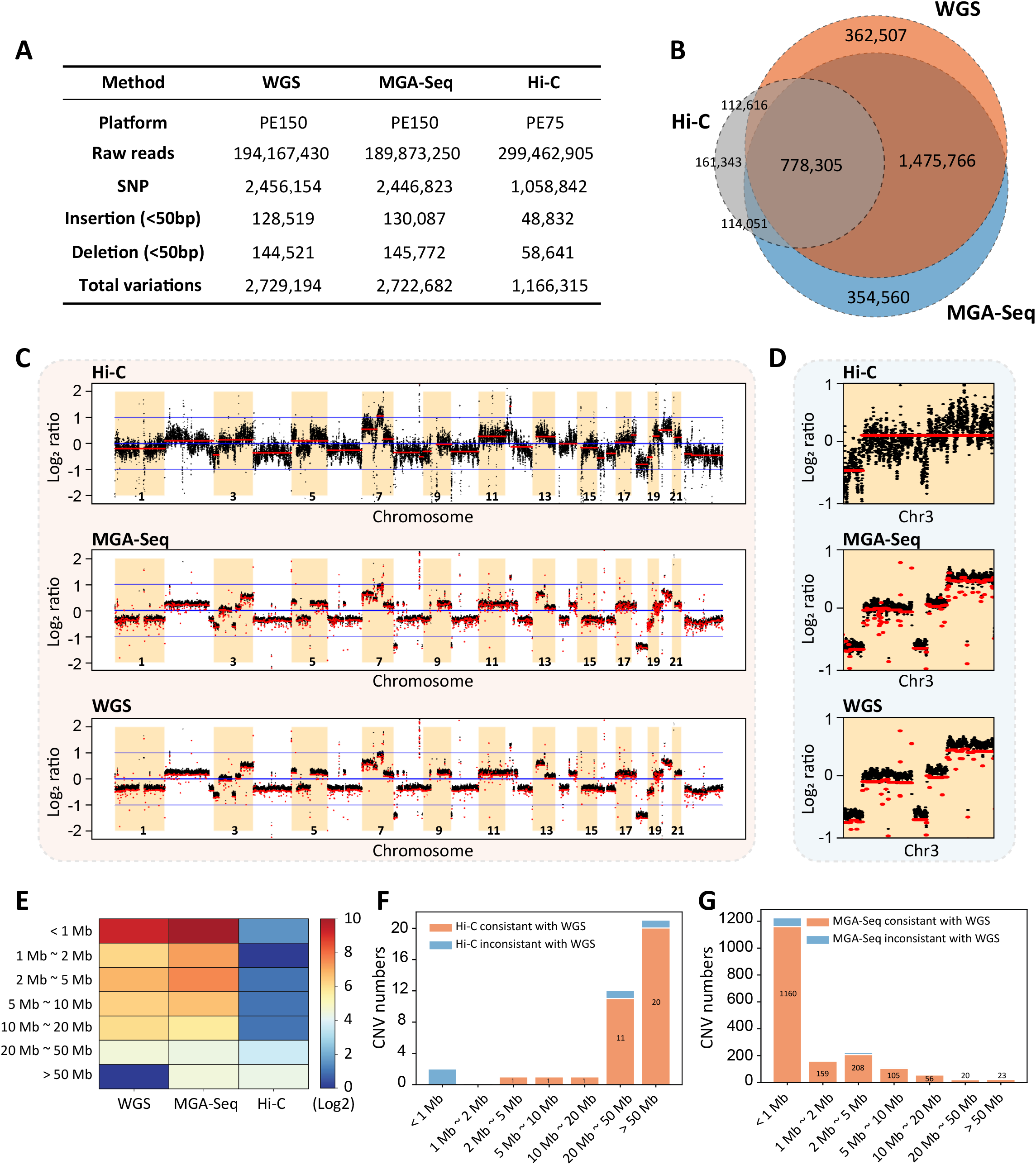
Detection of SNPs, indels, and CNVs by MGA-Seq. **(A)** Comparison of the numbers of SNPs and indels (< 50bp, include insertions and deletions) identified by WGS, MGA-Seq, and Hi-C. **(B)** Overlap of the SNPs and indels between MGA-Seq, WGS, and Hi-C. **(C)** Comparison of log_2_ copy ratios calculated using reads coverage between Hi-C, MGA-Seq, and WGS. **(D)** Comparison of the CNVs on chromosome 3 identified by Hi-C, MGA-Seq, and WGS. **(E)** Statistics of the number and size distribution of CNVs identified by Hi-C, MGA-Seq, and WGS. **(F)** Consistency of the CNV segments (categorized by size) detected by Hi-C and WGS. Overall, Hi-C cannot detect CNV with length less than 20 Mb. **(G)** Consistency of the CNV segments detected by MGA-Seq and WGS. The number and size distribution of CNV segments detected by MGA-Seq and WGS are highly consistent, especially for micro-CNVs (< 1Mb).

### Detection of chromosome copy number variation by MGA-Seq

To test the CNV detection capability of MGA-Seq, we plotted the log_2_ ratio of average read depths in 50 Kb bins across the genome, as shown in **Fig. 2C**. Our data showed that the genome coverage and uniformity of MGA-Seq are highly consistent with the gold standard WGS datasets and much higher than those of the Hi-C datasets. After zooming in on chromosome 3, we observed that Hi-C roughly divided chromosome 3 into two CNV intervals, whereas MGA-Seq accurately identified all the small copy number variation across the whole chromosome (**Fig. 2D**). Next, we systematically analysed the size and number of CNVs identified by these three methods (**Fig. 2, E-G**) and found that it was extremely difficult to detect CNVs less than 10 Mb by Hi-C (**Fig. 2, E and F**). In this scenario, the CNV detection capability of MGA-Seq is much better than that of Hi-C, especially for micro-CNVs (<1 Mb), which is highly consistent with WGS (**Fig. 2, E and G**).

### Identification of chromosomal translocations and breakpoints by MGA-Seq with single base-pair resolution

By using SW480 MGA-Seq sequencing datasets, we obtained the genome-wide chromosome contact matrix. As shown in **Fig. 3, A and B**, we identified 8 translocations and 1 inversion. Although MGA-Seq only used 190 million raw reads, the structural variants detected by MGA-Seq were completely consistent with *in situ* Hi-C with 300 million raw reads (**Fig. 3A**). To further identify the chromosomal translocation types and breakpoints of these translocations, we combined chromosome contact matrix, CNV, and split read information from MGA-Seq datasets and performed integrated analysis. Taking T(2;12)(q35;q12) as an example, from the CNV data, we observed that the copy number of chromosome 12 was increased, whereas the copy number of chromosome 2 was decreased downstream of the chromosome breakpoint (**Fig. 3C**), suggesting that unbalanced translocation occurred between chromosomes 2 and 12.

**Figure 3.**
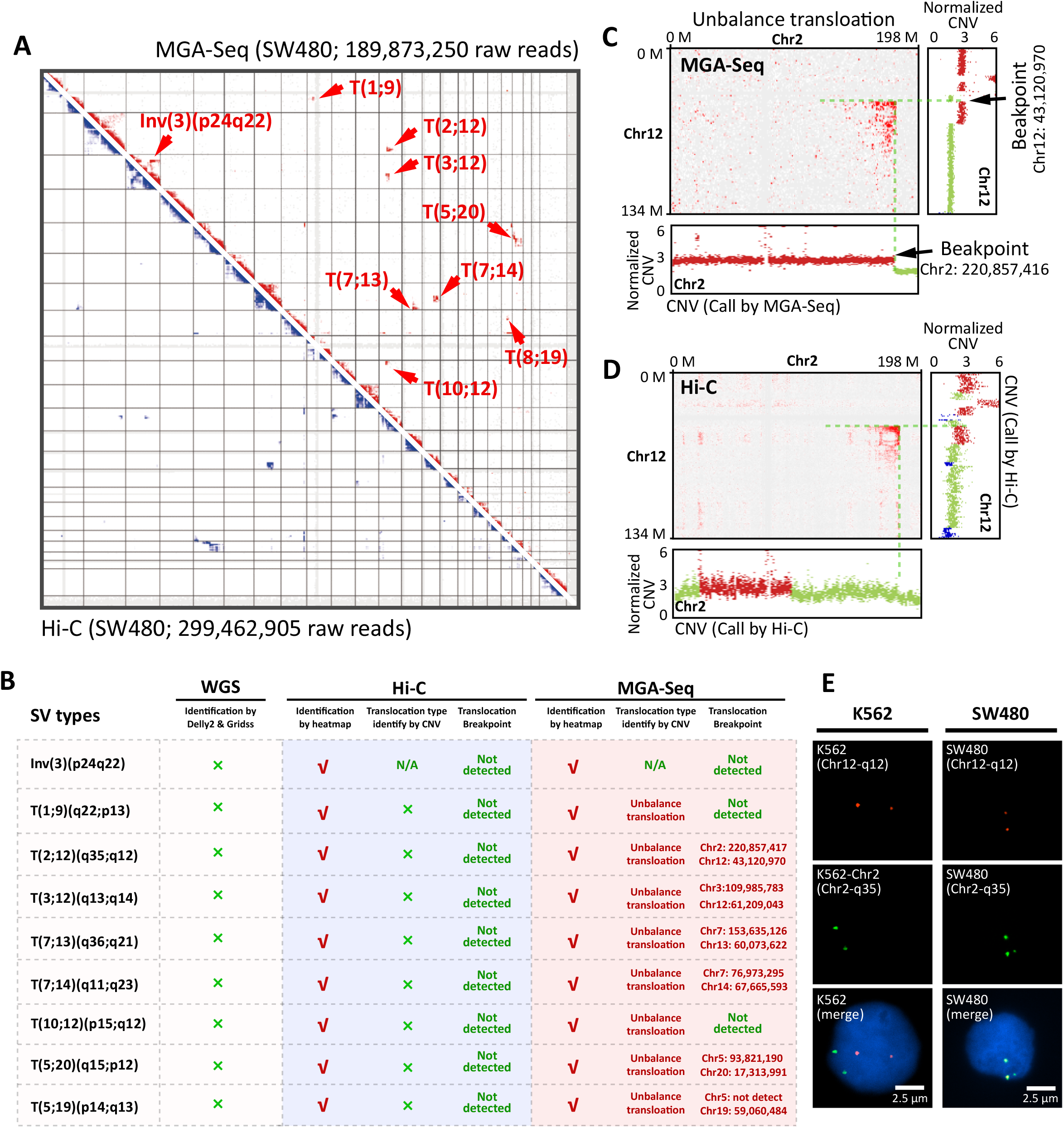
Identification of translocation types and breakpoints in SW480 at single base-pair resolution by MGA-Seq. **(A)** Dentification of translocation in the SW480 cell line by genomic contact matrix constructed with MGA-Seq datasets. The detected structural variations are indicated by arrows. **(B)** Translocation types and breakpoint information defined by MGA-Seq. **(C)** Application of integrated chromatin contact matrix, CNVs, and split reads analysis to identify translocation types and breakpoints between chr 2 and chr 12 at single base-pair resolution using MGA-Seq datasets. **(D)** Identification of translocation types and breakpoints between chr 2 and chr 12 using Hi-C datasets. **(E)** Validation of the T(2;12)(q35;q12) translocation in SW480 cells by DNA FISH. FISH probes for 12q12 and 2q35 were directly labeled with Alexa Fluor 555 (red) and Alexa Fluor 488 (green), respectively. K562 cells without the T(2;12)(q35;q12) translocation were used as a control.

Based on the split reads information in MGA-Seq, we further identified that the translocation breakpoint is located at chr2: 220,857,416 and chr12: 43,120,970 (**Fig. 3C**). In contrast, due to the low genome coverage and depth of Hi-C, it is not feasible to precisely determine the type and breakpoint of translocation (**Fig. 3, B and D**). In this scenario, MGA-Seq identified that all 8 chromosomal translocations in the SW480 cell line were unbalanced translocations. Notably, we were able to pinpoint the breakpoints of 6 out of 8 translocations sites at single-base resolution (75.0 %). We also used WGS data with the same sequencing depth as MGA-Seq to identify the translocations. As there is no chromosome interaction information in this dataset, none of the chromosomal translocations were found (**Fig. 3B**). Moreover, we verified the T(2;12)(q35;q12) translocation by two-colour DNA fluorescence *in situ* hybridization (FISH). As shown in **Fig. 3E**, chromosomes 2 and 12 were indeed fused together in SW480 cells, supporting the integrity of MGA-Seq.

Furthermore, to test the chromosomal translocation detection capability of MGA-Seq in clinical samples, we collected peripheral blood from two infertile patients with known translocation sites and constructed an MGA-Seq library. By combining the chromosome interaction matrix and CNV data, we detected a T(10;22)(p12;q13) translocation in sample 1 (**fig. S3A**) and a T(9;11)(q21;p14) translocation in sample 2 (**fig. S3B**), which are consistent with the known translocation sites identified by karyotyping. In addition, based on the split reads, we pinpointed the precise location of the breakpoints with single base-pair resolution (**fig. S3, A and B**). Next, we analysed the CNV information of these two samples based on the MGA-Seq data to determine the translocation type. Our data showed that there are no chromosome copy number changes around the translocation breakpoint, meaning that both infertile patients carry balanced translocations. Together, these data demonstrated that MGA-Seq can detect specific chromosomal translocation types and the corresponding breakpoint with high efficacy and low cost.

### Detection of ecDNA and HSR by MGA-Seq

There are two types of focal amplifications, extrachromosomal DNA (ecDNA) and intrachromosomal HSRs. Due to the high mobility and dramatic amplification amount of ecDNA, we speculated that ecDNA can randomly interact with each chromosome with a significantly higher interaction frequency than the normal interchromosome interaction, while HSRs only interact strongly within the specific chromosomes (**Fig. 4A**). To prove this hypothesis, we selected the ecDNA-positive cell line COLO320-DM (*7*) and the HSR-positive cell line SW480 (*38*) for MGA-Seq analysis. First, MYC amplifications in the form of ecDNA in COLO320-DM cells and in the form of HSR in SW480 cells were confirmed by DNA FISH (**Fig. 4, B and C**). In comparison to HSR-positive SW480 cells, ecDNA-positive COLO320-DM cells showed MYC amplification throughout the nucleus (**Fig. 4, D and E**). Furthermore, CNV analysis based on the MGA-Seq dataset accurately located the MYC amplification regions in these two cell lines (**Fig. 4, F-I**).

**Figure 4.**
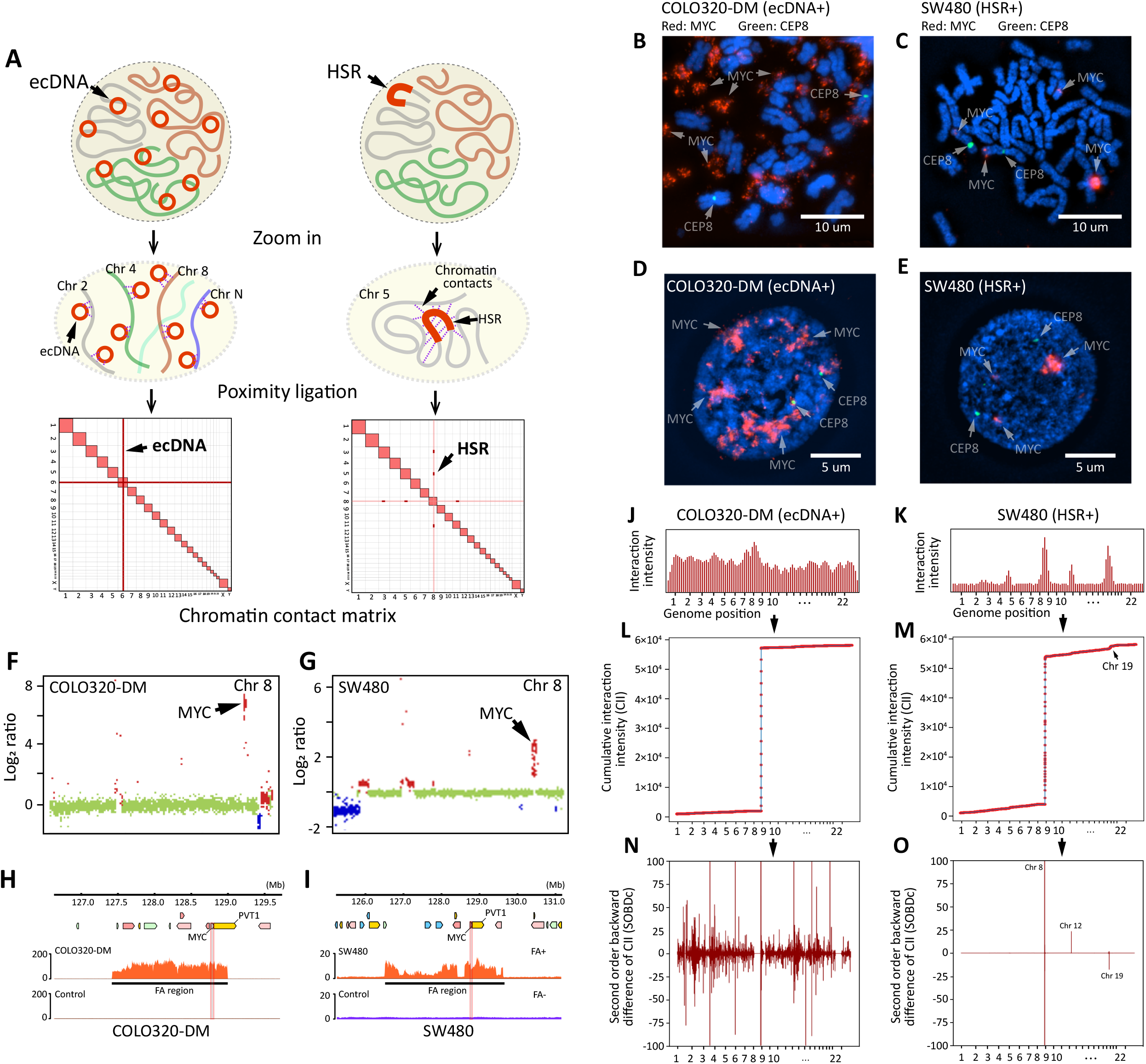
Identification of ecDNA and HSR by MGA-Seq. **(A)** Putative diagram of inter-chromosomal interaction pattern differences between ecDNA and HSR positive cell line. **(B-E)** Validation of *MYC* amplification in COLO320-DM and SW480 cell lines by DNA FISH. The red signal represents *MYC* and the green signal represents the centromere of chr 8. **(F and G)** Copy number variation analysis of chr 8 in COLO320-DM and SW480 cell lines. Gains and losses of copy number are shown in red and blue, respectively. **(H and I)** Location of the *MYC* amplification region in COLO320-DM and SW480 cell lines. **(J and K)** Interaction intensity between the focal amplified region and whole genome. **(L and M)** Cumulative interaction intensity curve of COLO320-DM and SW480 cell lines. The x-axis represents the genome position, 100 kb bin size. The Y axis represents the accumulation of interaction intensity. **(N and O)** Plotted the second order backward difference (SOBD) value across the genome of COLO320-DM and SW480 cell lines in 100-kb bin size.

Next, we constructed the chromatin interaction matrix using MGA-Seq data. Since the amplified ecDNAs were randomly distributed in the nucleus (**Fig. 4D**), the ecDNA fragments were unbiasedly ligated to all the chromatin fragments upon proximity ligation and thus presented a strip-like structure in the whole chromatin contact matrix (**Fig. S4A**). In contrast, as HSR is amplified on specific chromosomal regions (**Fig. 4, C and E**), it only shows strong interchromosomal interactions on certain chromosomes (**fig. S4B**), which is consistent with our hypothesis (**Fig. 4A**). In addition, we observed the same interchromosomal interaction pattern in ecDNA-positive cell lines TR14 and SUN16 (*7, 9*) (**fig. S4, C and D**). From the interchromosomal interaction matrix of SW480, we found that the MYC focal amplification region has a strong interaction with 19q13.3, indicating that MYC is likely to be amplified on chr19 (**fig. S4, E and F**). This finding is consistent with a previous report (*38*).

Since the judgement dependent on the naked eye is subjective and differs among individuals, we performed genome-wide interaction fluctuation analysis (GWIFA) on the focal amplification regions (**Fig. 4, J-O**) for more objective identification of HSR and ecDNA (see Methods). First, we divided the genome into fixed-size bins and calculated the interaction intensity between the amplified region and each bin (**Fig. 4, J and K**). The cumulative interaction intensity curve was then plotted as shown in **Fig. 4, L and M**. Next, second-order backwards difference (SOBD) analysis was applied to evaluate the fluctuation of the cumulative interaction intensity curve (**Fig. 4, N and O**). As HSR is amplified on the specific chromosome, the value of SOBD fluctuates dramatically at specific genomic locations (**Fig. 4O**). However, ecDNA has strong interactions with distinct strengths across the whole genome. Thus, the value of SOBD fluctuates greatly throughout the whole genome (**Fig. 4N, and fig. S4, G and H**).

### Delineation of the architecture of focal amplification in K562 cells

Our MGA-Seq analysis of K562 cells identified an abnormal increase in chromosome copy number on specific regions on chromosomes 9, 13, and 22 (**fig. S5A**). After zooming in on the abnormally amplified regions, we identified six precisely amplified subregions, one on chromosome 9, four on chromosome 13, and one on chromosome 22, which were named “A” to “F”, respectively (**Fig. 5A**). Based on genome-wide interaction fluctuation analysis (GWIFA), we found that these regions were amplified in K562 cells in the form of HSR rather than ecDNA (**fig. S5B**). Notably, we observed strong interactions between these amplified regions, suggesting that these regions are spatially close together, which likely originate from the same HSR (**Fig. 5B**). Taking the “B”, “C”, and “D” amplified regions of chromosome 13 as examples, these three regions are in high contact with each other and form a high-density topologically associating domain (TAD)-like structure (*39*) (**Fig. 5C**). Such abnormal genome amplification and TAD-like structures were absent in healthy human peripheral blood cells (**Fig. 5D**).

**Figure 5.**
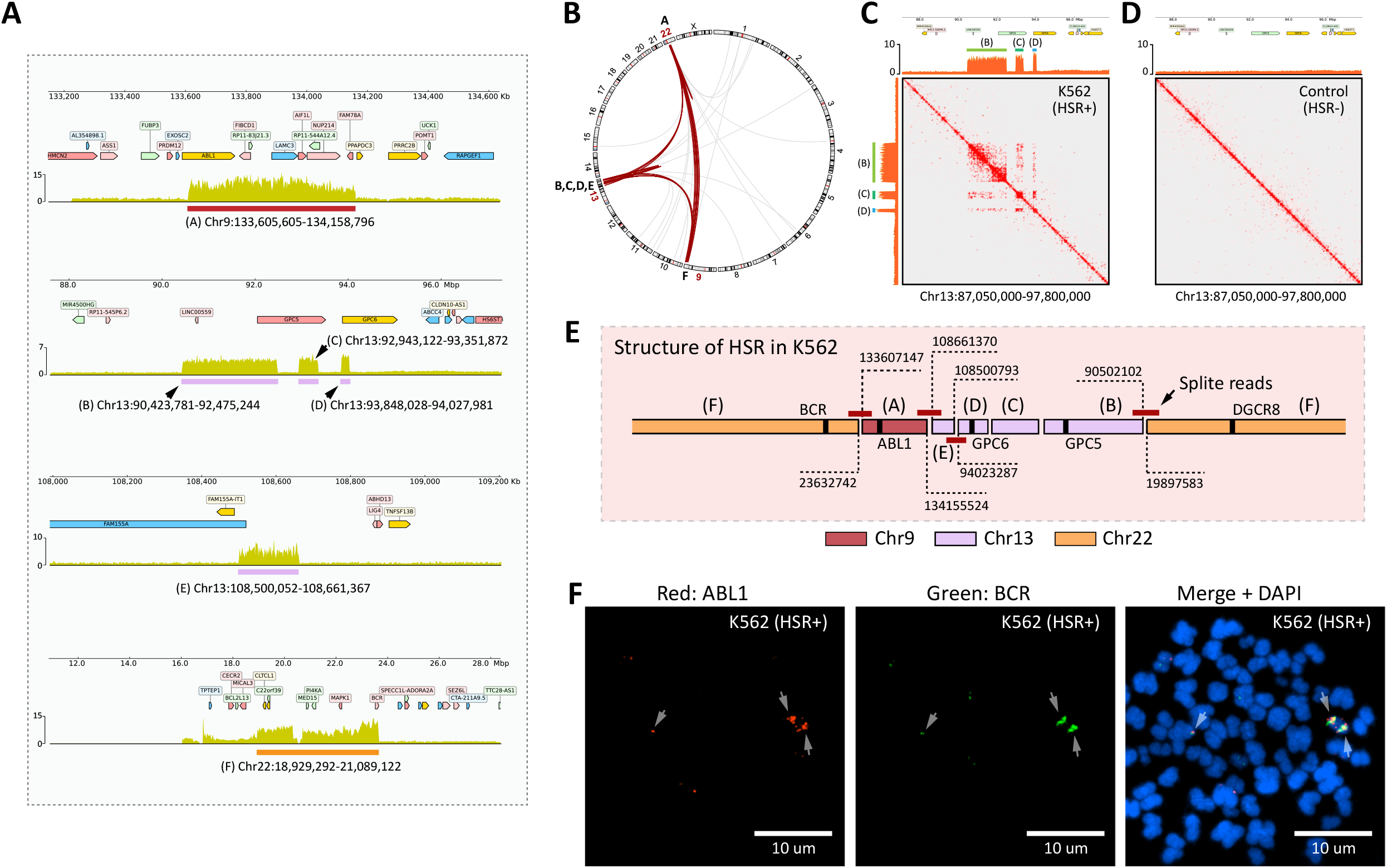
Deciphering the spatial structure of homogenously staining region (HSR) in K562 cell line. **(A)** Location of the amplification region on chr 9, 13, and 22. **(B)** Circos plots of the chromatin interactions mediated by amplification regions across all 23 chromosomes in K562 cell lines. The interactions between chromosomes 9, 13, and 22 are marked with red lines. **(C and D)** Comparison of chromatin contact matrix of amplification region (Chr13:90423781-92475244, Chr13:92943122-93351872, and Chr13:93848028-94027981) between K562 cell line and healthy human peripheral blood cells (control). **(E)** Assembling the amplified regions from “A” to “F” with split reads. The breakpoint of the amplification regions is marked in the figure. **(F)** Metaphase analysis and DNA FISH to validate the location of the *ABL1* amplification region and the *BCR* amplification region in K562 cell line. FISH probes for the *ABL1* amplification region and the *BCR* amplification region were directly labeled with Alexa Fluor 555 (red) and Alexa Fluor 488 (green), respectively.

Since the MGA-Seq dataset contains whole-genome sequencing information, we extracted the split reads located at the boundaries of these six amplified regions (table S1) and assembled the structure of HSR. In the K562 cell line, *ABL1* in the “A” amplification region, *GPC5* in the “B” amplification region, *GPC6* in the “D” amplification region, and *DGCR8* and *BCR* in the “F” amplification region were spliced to form a repeating HSR (**Fig. 5E and fig. S6**). To validate the HSR structure predicted by MGA-Seq, we compared our predicted results with published K562 third-generation sequencing data (*36*). Our analysis showed that *ABL1* in chromosome 9, *GPC5* and *GPC6* in chromosome 13, and *DGCR8* and *BCR* in chromosome 22 indeed come from the same scaffold, which is highly consistent with our results. Finally, we applied DNA FISH to verify the spatial location of the *ABL1* amplification region on Chr9 and the *BCR* amplification region on chromosome 22. As shown in **Fig. 5F**, *ABL1* and *BCR* indeed come from the same HSR.

### Identification of focal amplification in tumour tissue

Next, we applied MGA-Seq to tumour samples and detected 40 focal amplification regions in one renal cancer tissue (**fig. S**7 and **table S2**). The length distribution of these regions varies from 4.2 Kb to 2.53 Mb (**table S2**). These amplified regions contain a large number of immune genes, oncogenes, and enhancers, such as *CHD1L, BCL6, JAK2, PD-L1*, and *CDK4* (**Fig. 6A and table S2**). In addition, the RNA transcription level of these genes within the amplified region was significantly higher than that of the normal kidney tissue control (**Fig. 6A and fig. S8A**). Through MGA-Seq chromatin contact matrix and GWIFA (**fig. S8, B and C**), we identified that these FA regions are amplified in the form of HSR. Of note, these amplified regions are not independent but contact each other at the spatial level (**Fig. 6B**).

**Figure 6.**
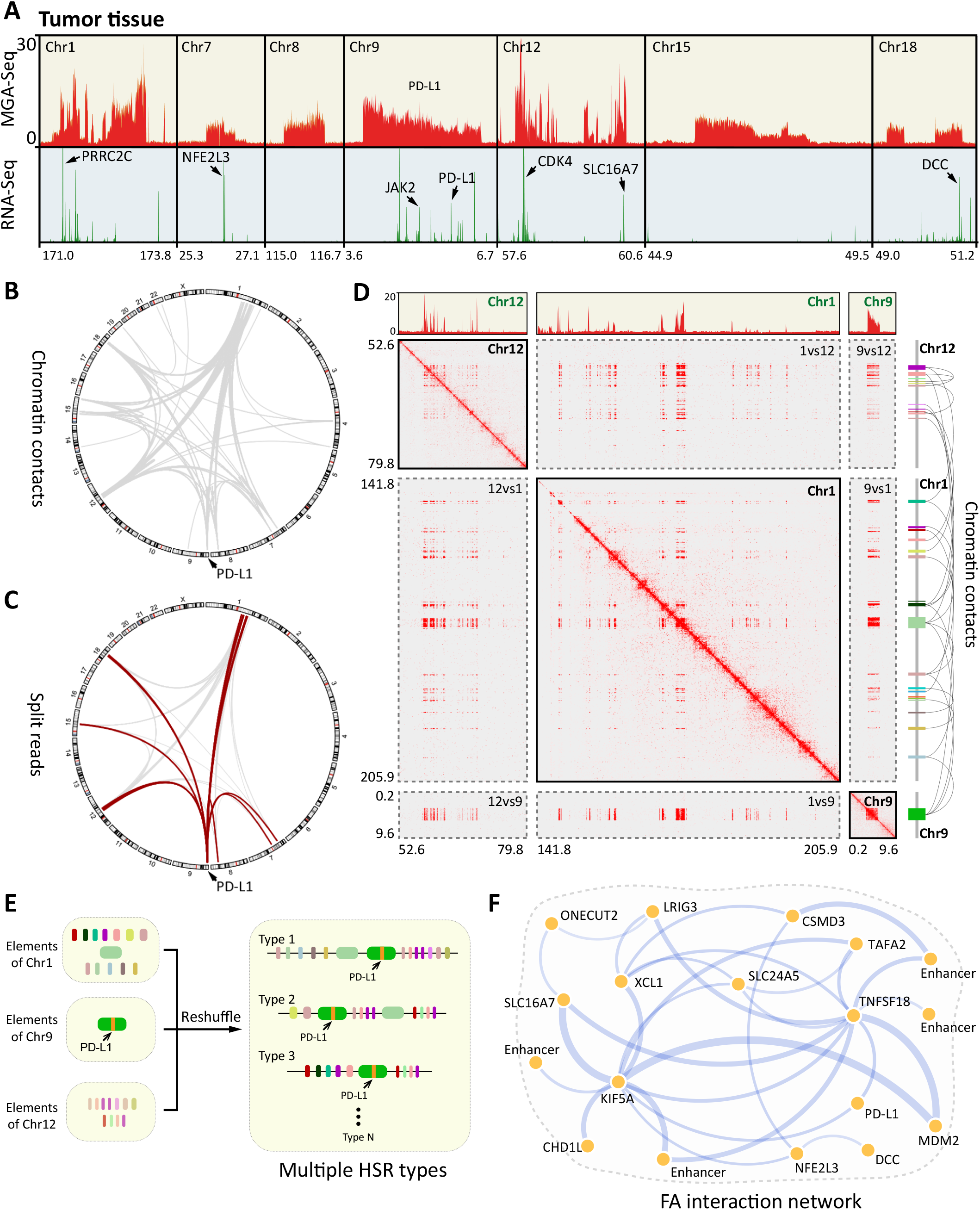
Heterogeneity of focal amplification in renal cancer tissue. **(A)** Sequencing reads coverage and RNA expression level in typical focal amplification regions of a renal cancer tissue sample. **(B)** Circos plots of the chromatin interactions mediated by focal amplification regions across all 23 chromosomes in renal cancer tissue. **(C)** Circos plots of the split reads mediated by focal amplification regions across all 23 chromosomes. The split reads aligned to the PD-L1 amplified region are marked with red lines. **(D)** Chromatin contact matrix between the amplified regions of chr1, chr9, and chr12, and sequencing reads coverage within these amplified regions. **(E)** Diverse structures of HSR in the renal cancer tissue sample. **(F)** Interaction network of amplified oncogenes in the renal cancer tissue sample. Different amplified oncogenes are assembled by split reads. The thickness of the line indicates the chromatin contact strength.

FA in tumour tissue is highly heterogeneous compared to single-cell-derived cell lines. Taking the PD-L1 amplification region on chromosome 9 in this tumour as an example, this region can be spliced with multiple FA regions, as indicated by the split reads in the MGA-Seq dataset, suggesting that multiple types of HSR coexist in this heterogeneous tumour tissue (**Fig. 6C**). To verify this result, we performed single-molecule nanopore sequencing on the same tumour sample, which revealed highly consistent inter- and intrachromosomal structural variation as with the MGA-Seq dataset (**fig. S8D**). The inter- and intrachromosomal interaction analysis of chr1, chr9, and chr12 amplification regions based on the MGA-Seq chromatin contact matrix (**Fig. 6D**) identified highly complicated and heterogeneous spatial architectures of these FA regions. For example, chr1 and chr12 show an “uneven amplification” pattern, meaning that in a certain chromosome interval, only some regions were amplified, such as the regions containing proto-oncogenes, immune genes, and some regulatory elements (**Fig. 6, A and D**). These genes and regulatory elements are spliced together and eventually form a variety of HSRs (**Fig. 6E**). Based on the chromatin contact information and split reads, we constructed an interaction network of these amplified oncogenes, in which the TNFSF18 region was connected with 11 amplified regions as supported by the split reads (**Fig. 6F**).

## Discussion

The occurrence of tumours, infertility, and rare diseases are closely related to focal amplification (*34*) and structural variation (*4, 5*). These genetic diseases affect hundreds of millions of people around the world and have become a major human health concern. An effective multiple genetic abnormality detection method is highly desired for clinical diagnosis. In this study, we developed multiple genetic abnormality sequencing (MGA-Seq) to simultaneously detect SNPs, CNVs, SVs, and the spatial architecture of FAs and distinguish ecDNA from HSR. Taking advantage of the versatility of reaction buffers, all the MGA-Seq library construction steps are carried out in a single tube, which can minimize sample loss due to buffer exchanges and simplify the operation. Notably, MGA-Seq takes only 9 hours and costs just 56 dollars to complete all the sequencing library construction steps. It demonstrated robust detection capability for both small (SNPs and INDELS) and large (CNV, SV, HSR, and ecDNA) genomic abnormalities and has great potential for clinical and scientific research applications.

ecDNA is prevalent in at least 30◻different cancer types, is closely associated with cancer progression (*11, 40*), and might be used as a potential prognostic marker. However, there is still a lack of an unbiased and efficient detection method in clinical practice. While AmpliconArchitect can be used for ecDNA prediction (*34*), the identification of ecDNA based on the WGS dataset generally has a high false-positive rate. For instance, in the cell line K562 in this study, due to the head-to-tail tandem duplication HSR structure (**Fig. 5E**), a large number of split reads also presented a circle junction-like structure. Circle-Seq can effectively analyse the structure of circular DNAs (*41, 42*). However, the DNA extraction process of this method can easily destroy the circular structure of ecDNA. Moreover, Circle-Seq is based on rolling-circle DNA amplification; it preferentially amplifies smaller circular DNAs, resulting in biased amplification results. Here, we demonstrated that MGA-Seq can unbiasedly detect the presence of ecDNA in both cell lines and clinical samples. Importantly, we proposed an ecDNA detection algorithm, GWIFA, and successfully differentiated ecDNA and HSR. Of note, MGA-Seq can reveal trans interactions between ecDNA and the genome and decode the ecDNA network, which could facilitate the exploration of the potential regulation of the expression inside of the ecDNA.

Chromosomal translocations can be divided into unbalanced and balanced translocations. Unbalanced translocation usually occurs with an altered chromosomal copy number at the breakpoint (gain or loss of genetic material), resulting in abnormal gene expression. A large number of unbalanced translocations have been found in cancer cells (*43, 44*), especially in blood tumour genomes (*45, 46*). Balanced translocations do not have any genetic material changes. These translocation carriers usually have normal phenotypes and intelligence but can produce various unbalanced rearranged gametes during germ cell meiosis, resulting in infertility, abortion, stillbirth, and multiple malformations (*6, 47, 48*). Thus far, it is still challenging to precisely identify the specific translocation types by a simple and cost-effective method. As MGA-Seq contains CNV and chromatin contact information, it can guide translocation breakpoint searching and facilitate to identifying translocation types and breakpoints. Here, we revealed the translocation types and breakpoints of infertile couples by MGA-Seq. With this important information, high-quality blastocysts can be quickly screened by PCR before blastocyst transfer during *in vitro* fertilization, which greatly reduces the cost and time of traditional whole genome sequencing for each blastocyst.

Together, we developed a simple, cost-effective and robust MGA-Seq to simultaneously detect SNPs, CNVs, SVs, and the spatial architecture of FA and distinguish ecDNA from HSR in a single tube experiment. We successfully identified small SNPs/INDELs and large genomic structural variations in clinical and cell line samples, decoded the focal amplification spatial structure in SW480, COLO320-DM, and K562 cell lines, and constructed interaction networks of the amplified proto-oncogenes in a clinical kidney cancer tissue sample. Our data revealed that focal amplification is highly diverse in tumour tissue compared to single-cell-derived cancer cell lines. In the future, it would be important to develop single-cell MGA-Seq for diverse ecDNA detection in single cells or highly heterogeneous cancer cells. With its multifunctional and cost-effective advantages, we expect MGA-Seq to be extensively applied for the diagnosis of cancer, infertility, and rare diseases and may greatly facilitate the investigation of the genomic mechanism for genetic diseases.

---

**Figure S1. Flow-chart of MGA-Seq data analysis**. After sequencing, all sequencing reads were used to generate chromatin contact matrix by juicer pipeline. The reads without proximity ligation junction were used to detect small indels and SNPs (< 50bp), CNVs, split reads, and genomic amplification regions. With the integrated analysis of chromatin contact matrix, these datasets can be used to decode the type and breakpoints of translocations, distinguish ecDNA from HSR, predict the focal amplification structure, and construct FA region interaction network.

**Figure S2. Comparison of the sequencing depth and coverage of MGA-Seq, WGS, and Hi-C. (A)** Scatter plot of sequencing depth and coverage for each chromosome. Blue points represent MGA-Seq, yellow points represent MGA-Seq, green points represent Hi-C. X-axis represents coverage, and Y-axis represents sequencing depth. **(B)** Histogram of coverage for each chromosome. Blue represents MGA-Seq, yellow represents MGA-Seq, and green represents Hi-C. **(C)** Histogram of sequencing depth for each chromosome. Blue represents MGA-Seq, yellow represents MGA-Seq, and green represents Hi-C.

**Figure S3. Identification of translocation types and breakpoints by MGA-Seq. (A)** Identification of balance translocation T(10;22)(p12;q13) and genome breakpoint in patient 1. **(B)** Identification of balance translocation T(9;11)(q21;p14) and genome breakpoint in patient 2.

**Figure S4. Chromatin contact matrix and genome-wide interaction fluctuation analysis (GWIFA) of ecDNA-positive cell lines. (A-D)** Genome-wide chromatin contact matrix of COLO320-DM, SW480, TR14, and SUN16 cell lines. The amplified regions are marked with arrows. **(E)** The chromatin interaction matrix of SW480 cell line between chr 8 and chr 19. The *MYC* amplified region is marked with a dashed line in the figure. **(F)** MYC is amplified in the form of HSR on chr 19. **(G and H)** The second order backward difference (SOBD) value across the genome of TR14 and SUN16 cell lines in 100-kb bin size.

**Figure S5. Copy number variation (CNV) analysis of K562 cell line. (A)** CNV analysis of chromosomes 9, 13, and 22 in K562 cell line. Gains and losses of copy number are shown in red and blue, respectively. Representative genes located in amplification region are marked with arrows. **(B)** The second order backward difference (SOBD) value across the genome of K562 cell line in 100-kb bin size.

**Figure S6. Sequence and breakpoints of split reads used to assemble HSR in K562 cells**.

**Figure S7. CNV analysis of renal cancer tissue**. CNV analysis of chromosomes with abnormal amplification in renal cancer tissue. Gains and losses of copy number are shown in red and blue, respectively. Representative genes located in amplification region are marked with arrows.

**Figure S8. Verification of inter and intra chromosomal interaction between focal amplification regions in renal cancer tissue by nanopore. (A)** Volcano plots of differential expression genes between renal cancer tissue and normal kidney tissue control. **(B)** Genome-wide chromatin contact matrix of renal cancer tissue. Potential HSR regions are marked with arrows in the figure. The inter-chromosomal contacts between the focal amplification regions and Chr1 and Chr12 are zoomed in. **(C)** The second order backward difference (SOBD) value across the genome of the renal cancer tissue in 100-kb bin size. **(D)** Validation of split reads and chromatin interactions across focal amplification regions with Nanopore long reads.

**Figure S9. Distribution of fluctuation score (FS) in different cell lines**. Blue bars indicate HSR-positive cell lines and yellow bars indicate ecDNA-positive cell lines. The Y axis represents the value of FS.

**Table S1. The split reads located at the boundaries of focally amplified regions in K562 cells**.

**Table S2. The regions of focal amplification in the renal cancer tissue.**

## METHODS

### MGA-Seq library construction

#### 1. Preparation of cell suspension

For tumor tissue, 0.5 cm^3^ tissue blocks were used and minced through a 40 μm strainer to obtain single cell suspension. For blood samples, we directly took 1 ml of anticoagulated whole blood, and centrifuged at 1500 g/min for 10 min to collect blood cells.

#### 2. Nuclei preparation

Cells were cross-linked with 0.5 % formaldehyde (Sigma) for 10 mins. The cross-linking reaction was terminated by glycine at a final concentration of 200 mM and lysed in lysis buffer (PBS contain 0.2% SDS) at room temperature for 5 min. After incubation, the nuclei were pelleted by centrifugation at 2,000 g/min for 5 min. The nuclei were transferred to 1.5 ml tubes and washed twice with PBS.

#### 3. *In situ* digestion

For *in situ* restriction enzyme digestion, 140 μl of ddH_2_O, 20 μl of 10% Triton X-100, 20 μl of 10× NEBuffer 2.1, and 20 μl of HindIII (NEB, 20 units/μl) were added to the nuclei pellet and digested for 1.5 h at 37 °C in thermomixer (Eppendorf) with rotation at 1000 r.p.m.

#### 4. End filling-in

Add 5 μl of dNTP mix (10 mM each) and 5 μl of DNA polymerase I Klenow fragment (NEB, M0210) to the reaction system, place the sample in thermomixer with rotation at 37 °C at 1000 r.p.m for 30 mins.

#### 5. *In situ* proximity ligation

Add 27.5 μl of H_2_O, 3 μl of ATP (adenosine-triphosphate, 10mM), and 10 μl of T4 DNA ligase (Thermo, EL0011) to the reaction system, and placed the tube on the rotating mixers for 2 h at room temperature with rotation at 20 r.p.m.

#### 6. Reversal of cross-linking and DNA purification

Add 20 μl of proteinase K (20 μg/ml) to the proximity ligation system, and then incubate at 60 °C for 2 hours. After digestion, the DNA was directly extracted using PCR Purification Kits (Zymo, D4013).

#### 7. Sequencing library construction

DNA sequencing libraries were prepared using the VAHTS Universal Plus DNA Library Prep Kit (NDM627) according to the manufacturer’s protocol.

### Metaphase analysis and DNA fluorescence in situ hybridization (FISH) assay

SW480 and COLO320-DM cell lines were treated with colchicine at final concentration 8 μg/ml for 24 hours. After cultivation, cells were collected by centrifugation at 1000 g/min for 10 minutes. Next, 10 ml of hypotonic KCl solution (0.075 M) were added to the cell pellet to resuspend the cells. After 30 min incubation at 37 °C, 2 ml of fixative (3:1 methanol:glacial acetic acid) were added to the cell suspension. The cell pellet was re-collected by centrifugation at 1000 g/min for 10 min and then resuspend in 5 ml of fixative (3:1 methanol:glacial acetic acid). After 5 min incubation, the cell pellet was re-collected by centrifugation at 1000 g/min for 10 minutes and resuspend in 1 ml of fixative. After fixation, 10 μl of the suspension were dropped on the glass slide and incubated in the prewarmed 2x SSC at 60 ◻ for 30 min. The cells were dehydrated sequentially in 70%, 85%, 100% ethanol solution. After ethanol dehydration, the cells were heated on a hot plate at 82 °C for 10 min in 80 % formamide (Sigma) and 2×SSC for DNA denaturation. Next, cells were incubated for 12 hours in hybridization solution with 2 μM DNA probes (MYC and CEP8, Spatial FISH Co. Ltd.) in the presence of 50 % formamide, 8% dextran sulfate sodium salt (Sigma), and 2× SSC. After hybridization, the cells were washing for three times with 20 % formamide and 3 times with 2×SSC. Finally, the slides were stained with DAPI (Life Technologies) and observed under super-resolution microscope (Nikon, N-SIM).

### RNA-Seq library preparation

RNA was extracted using the RNAiso Plus (Takara, 9109) according to the manufacturer’s protocol. Sequencing libraries were prepared using the VAHTS Stranded mRNA-Seq Library Prep Kit (Vazyme, NR602-02) according to the manufacturer’s protocol.

### Identification of SNP, indel, split reads, and CNV using MGA-Seq datasets

#### 1. Pre-analysis

FastQC (version: 0.11.5) (*49*) was used to assess the quality of raw reads. FASTP (version: 0.23.2) (*50*) was used to filter out the low-quality bases and adapter sequences. The clean read pairs which contained proximity ligation junction sequences “AAGCTAGCTT” were filtered out by Linux command line utility “grep”. The remaining reads were used for SNPs, indels, split reads, and CNV calling.

#### 2. SNP and indel calling

The remaining reads were aligned to the reference genome (hg19) and generated BAM file using BWA-MEM (version 0.7.17) (*51*). The BAM file was sorted by SAMtools (version 1.15.1) (*52*) and deduplicated by Sambamba (*53*) (version 0.6.6). Next, we used BaseRecalibrator (GATK, version 4.2.2) (*54*) to calibrate the base quality scores, and HaplotypeCaller (GATK, version 4.2.2) to detect SNPs and indels.

#### 3. Split reads calling

The deduplicated BAM file generated in SNP and indel calling step were used to identify split reads. The split alignment reads were extracted by SAMtools (version 1.15.1) with command line “samtools view test_deduplicated.bam | grep SA > test_split_reads.txt”.

#### 4. CNV calling

BIC-seq2(*55*) was used to derive CNV segments from reads coverage data. For more details, refer to the software manual “http://www.compbio.med.harvard.edu/BIC-seq/”. For the segmentation step, parameters were designed as binsize◻=◻50,000 bp and λ◻=◻2 to determine the final CNV breakpoints.

### Construction of genome-wide chromatin interaction matrix using MGA-Seq datasets

FastQC (version: 0.11.5) was used to assess the quality of raw reads. FASTP (version: 0.23.2) was used to filter out the low-quality bases and adapter sequences. All the remaining read pairs were used to generate the chromatin contacts matrix file (.hic) using Juicer software(*56*). For more details, refer to the software manual “https://github.com/aidenlab/juicer”.

### Identification of translocations types and breakpoints using MGA-Seq datasets

The chromatin contacts matrix file (.hic) was imported into Juicerbox (version: 1.9.8, https://github.com/aidenlab/Juicebox) software for visualization. The translocations and large structural variations were identified according to the inter-/intra-chromosome interaction patterns (*15, 57*). The types and breakpoints of translocations were identified according to the split reads and CNV information. For unbalanced translocations, the chromosomal copy number at the breakpoint were usually altered, while balanced translocations do not have any chromosomal copy number changes.

### Identification of SNP and indel using *in situ* Hi-C datasets

The *in situ* Hi-C datasets of SW480 cell line (*37*) were downloaded from Gene Expression Omnibus (GEO Accession: GSM3930294 and GSM3930295). The Hi-C ligation junction sequence “GATCGATC” and bases behind ligation junction were removed by FASTP (version: 0.23.2). An example command line is as follows:

1. fastp -i insitu_sw480_1.fq -o trim_sw480_1.fq -w 15 --adapter_sequence GATCGATC
2. fastp -i insitu_sw480_2.fq -o trim_sw480_2.fq -w 15 --adapter_sequence GATCGATC

Trimmed reads1 and reads2 were merged together by command line “cat trim_sw480_1.fq trim_sw480_2.fq > sw480_1_2.fq”. The merged reads file was used to identify SNPs and indels using the same parameters as MGA-Seq.

### Identification of CNV and translocation by *in situ* Hi-C datasets

All the raw Hi-C read pairs were used to detect CNVs. FastQC (version: 0.11.5) was used to assess the quality of raw reads, and FASTP (version: 0.23.2) was used to filter out the low-quality bases and adapter sequences. The CNV calling was carried out by BIC-seq2 (*55*). The observed values were the residuals from GAM Poisson regression, and the expected values were set to zero. Translocation detection was performed by HINT-TL as implemented in HINT (*58*), a computational method for detecting CNVs and translocations based on Hi-C data.

### Identification of SNP, indel, and translocations using WGS datasets

FastQC (version: 0.11.5) was used to evaluate the quality of raw reads. FASTP (version: 0.23.2) was used to filter out the low-quality bases and adapter sequences. The trimmed reads pairs were used to identify SNPs and indels. The parameters are exactly the same as MGA-Seq.

Structural variation identification were carried out using Delly2 (*59*) (version: 0.8.6) and Gridss (*60*) (version: 2.12.2) with default parameters. One WGS data from healthy person was served as control. Translocations that passed the internal quality control were merged with SURVIVOR (version: 1.0.7, parameters: 1000 1 1 1 0 30) (*61*). Only translocations supported by at least one definite split alignment read were retained.

### Genome-wide interaction fluctuation analysis (GWIFA)

According to the inter-chromosomal interaction feature of ecDNA and HSR, we designed a genome-wide interaction fluctuation analysis (GWIFA) to further characterize the inter-chromosomal interaction fluctuation of the focal amplification regions and defined a fluctuation score (FS) to distinguished ecDNA from HSR.

Firstly, we divided genome into fixed-sized bins (100 kb), and calculated the cumulative interaction intensity (CII) between the focal amplified regions and the whole genome (**Fig. 4, J-M**).

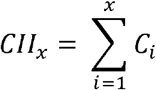

In the formula, x represents the genome position measured by the number of bin, and C_i_ represents the number of contact counts inside the ith bin. We recommend a linear fit on CII, which can eliminate the abnormal fluctuations caused by uneven sequencing.

Next, second order backward difference (SOBD) was introduced to further characterize the fluctuation of interactions across the genome (**Fig. 4, N and O**). Denoting OBD of CII as OBDc.

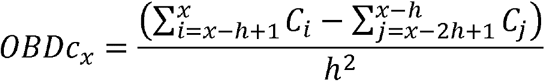

In the formula, h is a customizable space, default is 3.

We then defined a fluctuation score (FS) to distinguished ecDNA from HSR.

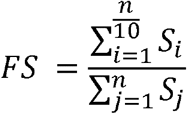

In the formula, S is descending sorted distribution of |OBDc| (absolute value of OBDc), n is the quantity of OBDc, T is a customizable parameter (T<1). After multiple rounds of testing, our suggested T is 0.6 (**fig. S9**).

ecDNA and HSR can be distinguished as follow:

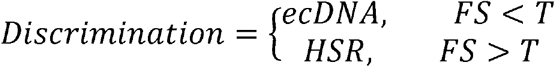

The complete analysis pipeline is available in: https://github.com/yanyanzou0721/GWIFA.

### Long-read sequencing (Nanopore) data analysis

The nanopore sequencing reads with quality score more than 7 were mapped to the reference genome hg19 using minimap2 (version: 2.17, -ax map-ont) (*62*). Structural variants were called using NanoSV(*63*) (version: 1.2.4) with default parameters. Only SV supported by at least one definite split alignment read were retained for subsequent statistics.

### RNA-seq data analysis

FastQC (version: 0.11.5) was used to assess the quality of the raw reads. FASTP (version: 0.23.2) was used to filter out the low-quality bases and adapter sequences. The clean reads were aligned to the hg19 using BWA-MEM (version: 0.7.17) with default parameters and sorted by Samtools (version: 1.15.1). Gene expression levels were assessed using featureCounts (version: 2.0.0) (*64*). Differential gene expression analysis was performed using DEseq2 (version: 1.20.0) (*65*). The complete analysis pipeline is available in https://github.com/GangCaoLab/NGS-pipelines/tree/master/RNA-Seq.

## Data availability

Data have been deposited in the Gene Expression Omnibus (GEO). To review GEO accession GSE205293, go to https://www.ncbi.nlm.nih.gov/geo/query/acc.cgi?acc=GSE205293, and enter token “epgtysmkzvuftmp” into the box.

