## Supplemental Fig 1 for "Robust identification of extrachromosomal DNA and genetic variants using multiple genetic abnormality sequencing (MGA-Seq)"

### Figure S1

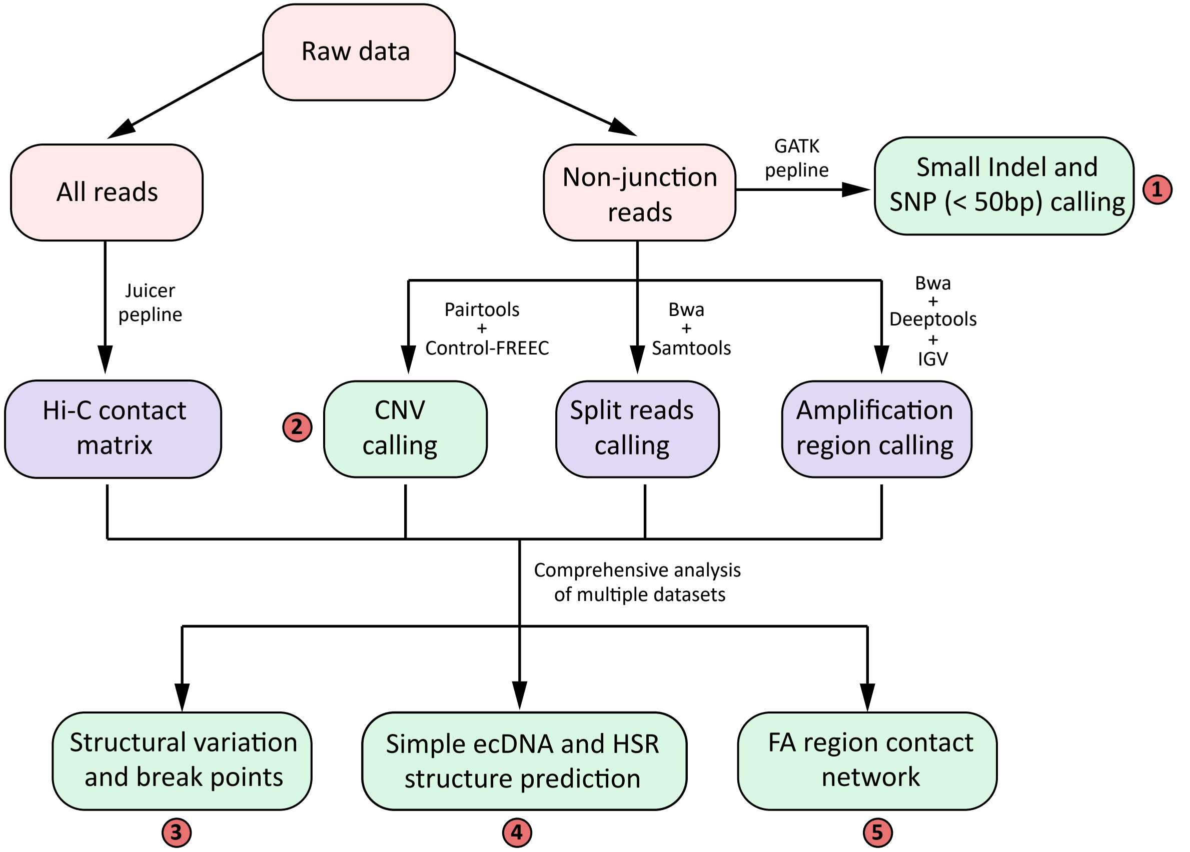


**Figure S1. Flow-chart of MGA-Seq data analysis.** After sequencing, all sequencing reads were used to generate chromatin contact matrix by juicer pipeline. The reads without proximity ligation junction were used to detect small indels and SNPs (< 50bp), CNVs, split reads, and genomic amplification regions. With the integrated analysis of chromatin contact matrix, these datasets can be used to decode the type and breakpoints of translocations, distinguish ecDNA from HSR, predict the focal amplification structure, and construct FA region interaction network.
