## Supplemental Fig 2 for "Robust identification of extrachromosomal DNA and genetic variants using multiple genetic abnormality sequencing (MGA-Seq)"

### Figure S2

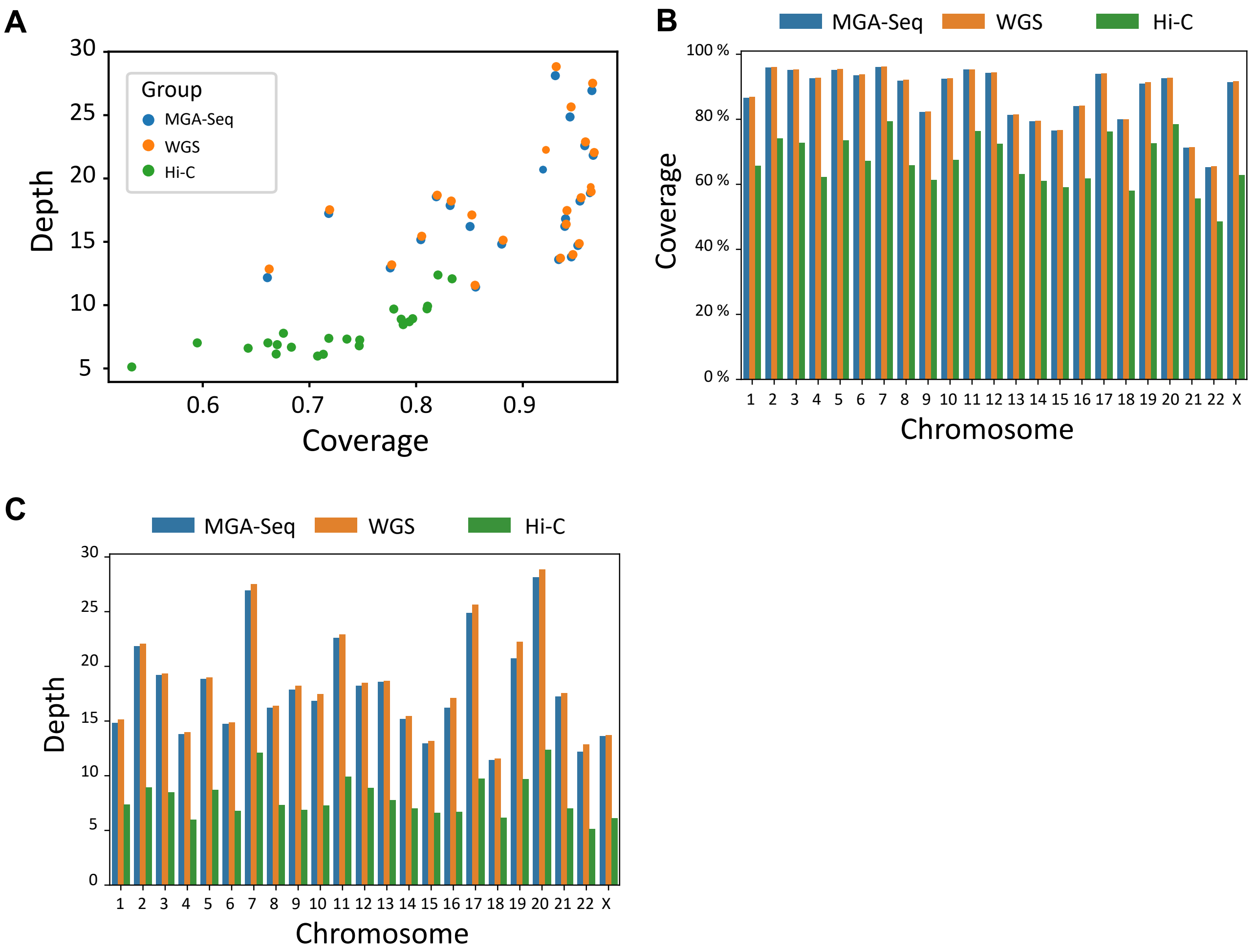


**Figure S2. Comparison of the sequencing depth and coverage of MGA-Seq, WGS, and Hi-C. (A)** Scatter plot of sequencing depth and coverage for each chromosome. Blue points represent MGA-Seq, yellow points represent MGA-Seq, green points represent Hi-C. X-axis represents coverage, and Y-axis represents sequencing depth. **(B)** Histogram of coverage for each chromosome. Blue represents MGA-Seq, yellow represents MGA-Seq, and green represents Hi-C. **(C)** Histogram of sequencing depth for each chromosome. Blue represents MGA-Seq, yellow represents MGA-Seq, and green represents Hi-C.
