## Supplemental Fig 3 for "Robust identification of extrachromosomal DNA and genetic variants using multiple genetic abnormality sequencing (MGA-Seq)"

### Figure S3

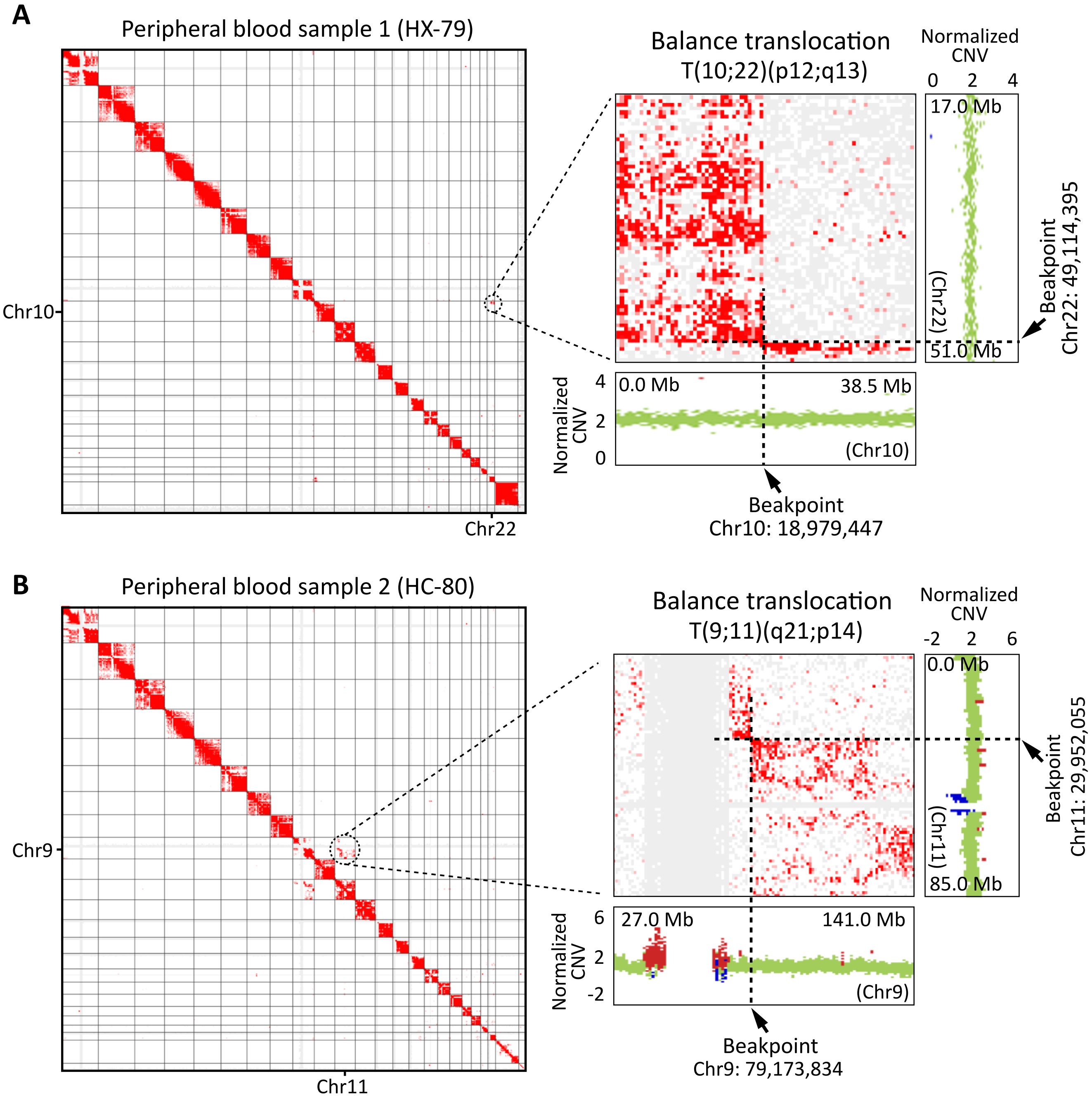


**Figure S3. Identification of translocation types and breakpoints by MGA-Seq. (A)** Identification of balance translocation T(10;22)(p12;q13) and genome breakpoint in patient 1. **(B)** Identification of balance translocation T(9;11)(q21;p14) and genome breakpoint in patient 2.
