## Supplemental Fig 4 for "Robust identification of extrachromosomal DNA and genetic variants using multiple genetic abnormality sequencing (MGA-Seq)"

### Figure S4

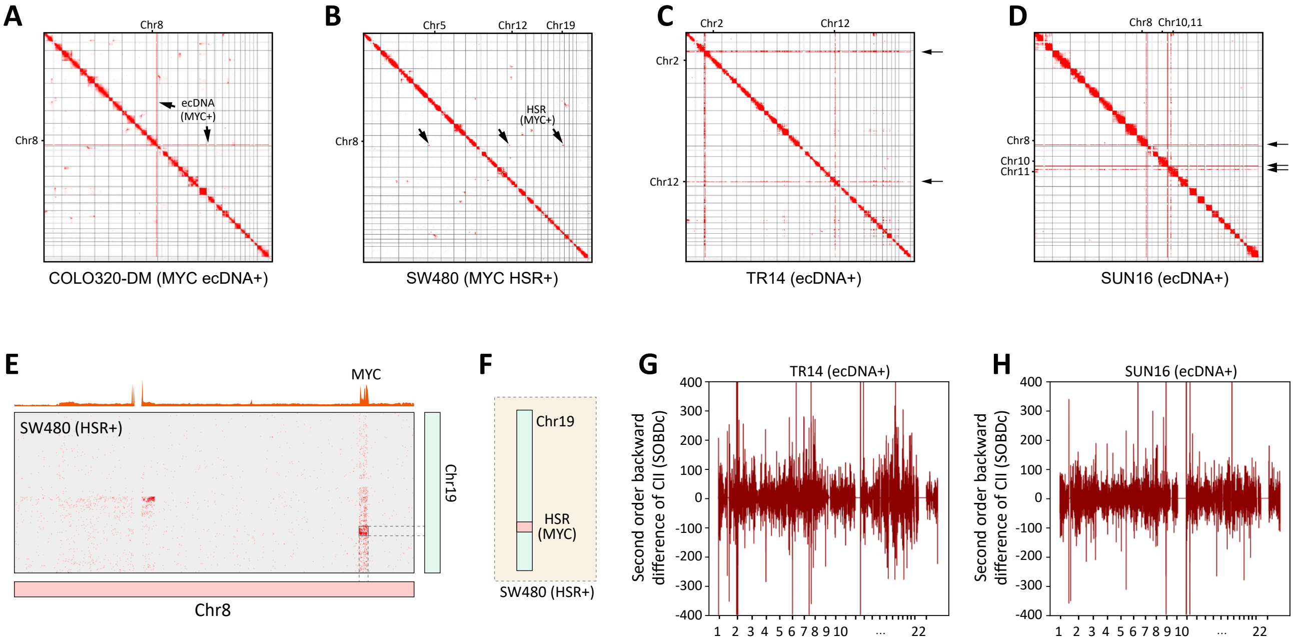


**Figure S4. Chromatin contact matrix and genome-wide interaction fluctuation analysis (GWIFA) of ecDNA-positive cell lines. (A-D)** Genome-wide chromatin contact matrix of COLO320-DM, SW480, TR14, and SUN16 cell lines. The amplified regions are marked with arrows. **(E)** The chromatin interaction matrix of SW480 cell line between chr 8 and chr 19. The MYC amplified region is marked with a dashed line in the figure. **(F)** MYC is amplified in the form of HSR on chr 19. **(G and H)** The second order backward difference (SOBD) value across the genome of TR14 and SUN16 cell lines in 100-kb bin size.
