## Supplemental Fig 5 for "Robust identification of extrachromosomal DNA and genetic variants using multiple genetic abnormality sequencing (MGA-Seq)"

### Figure S5

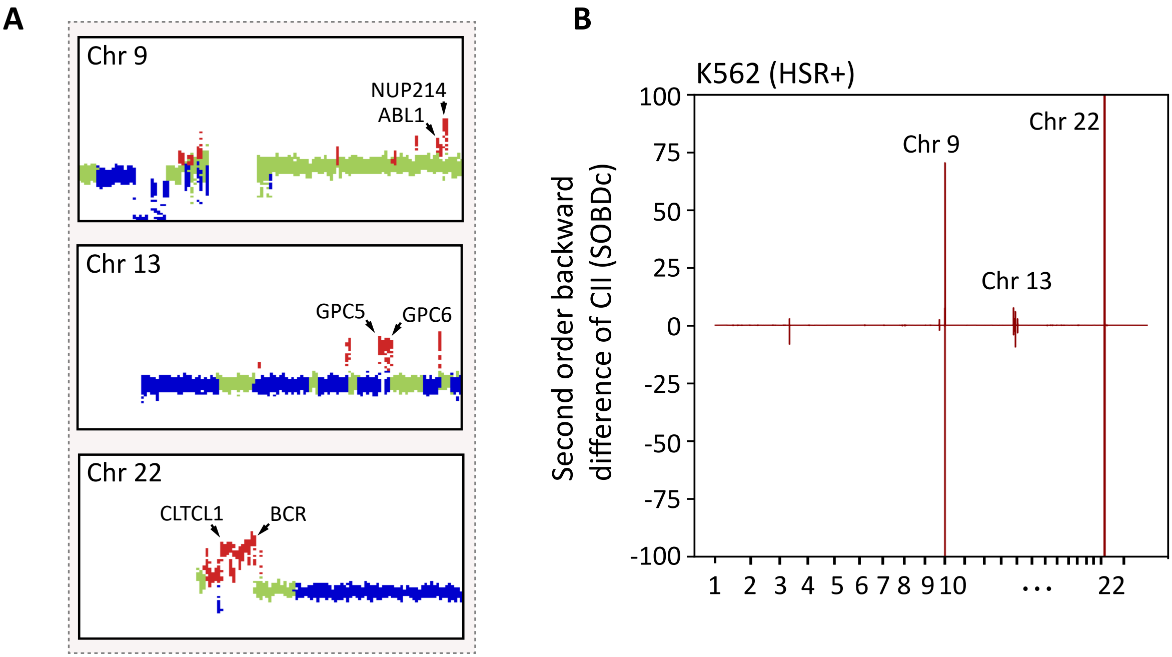


**Figure S5. Copy number variation (CNV) analysis of K562 cell line. (A)** CNV analysis of chromosomes 9, 13, and 22 in K562 cell line. Gains and losses of copy number are shown in red and blue, respectively. Representative genes located in amplification region are marked with arrows. **(B)** The second order backward difference (SOBD) value across the genome of K562 cell line in 100-kb bin size.
