## Supplementary figures and images for "Robust identification of extrachromosomal DNA and genetic variants using multiple genetic abnormality sequencing (MGA-Seq)"

### Supplemental Fig 6

#### Figure S6

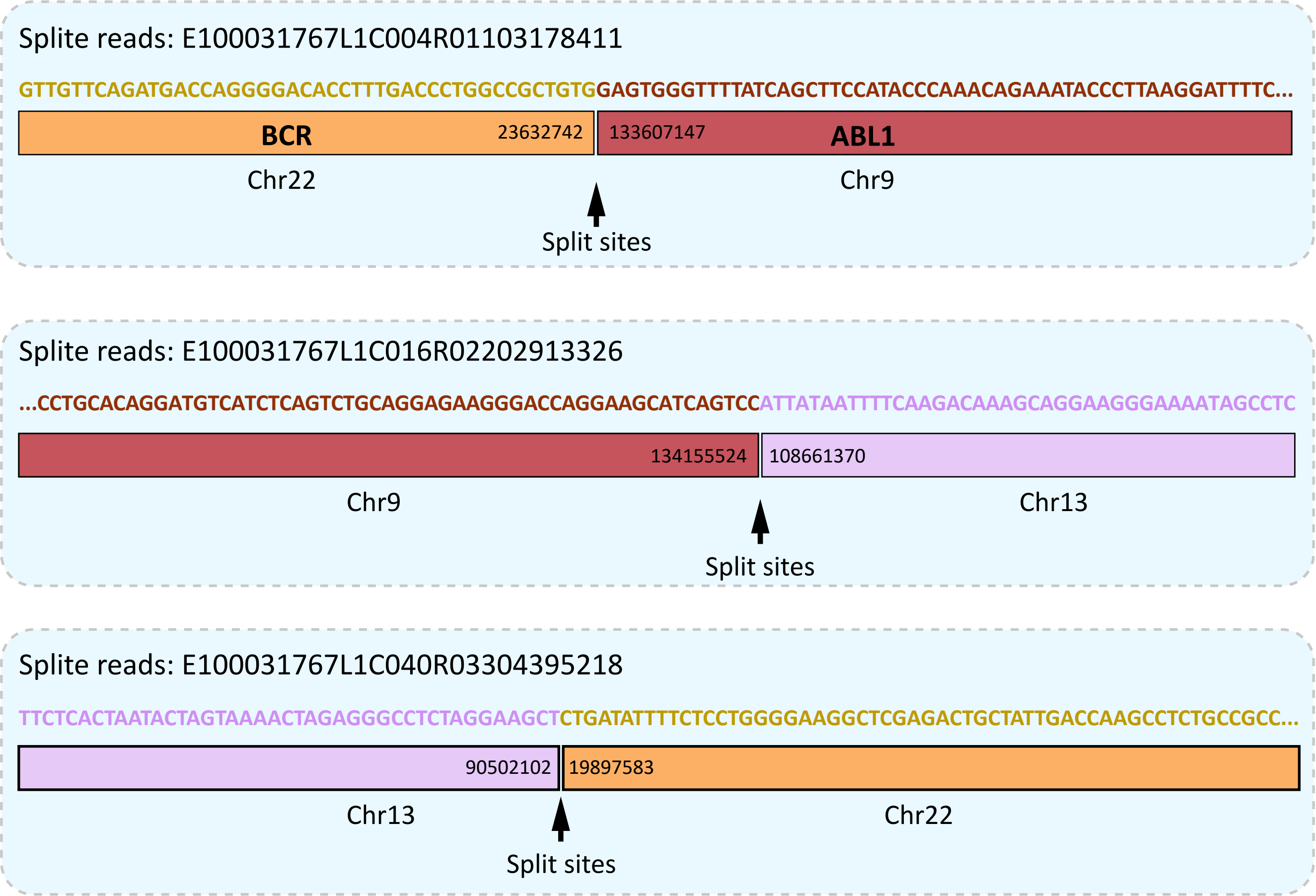


**Figure S6. Sequence and breakpoints of split reads used to assemble HSR in K562 cells.**
