## Supplemental Fig 7 for "Robust identification of extrachromosomal DNA and genetic variants using multiple genetic abnormality sequencing (MGA-Seq)"

### Figure S7

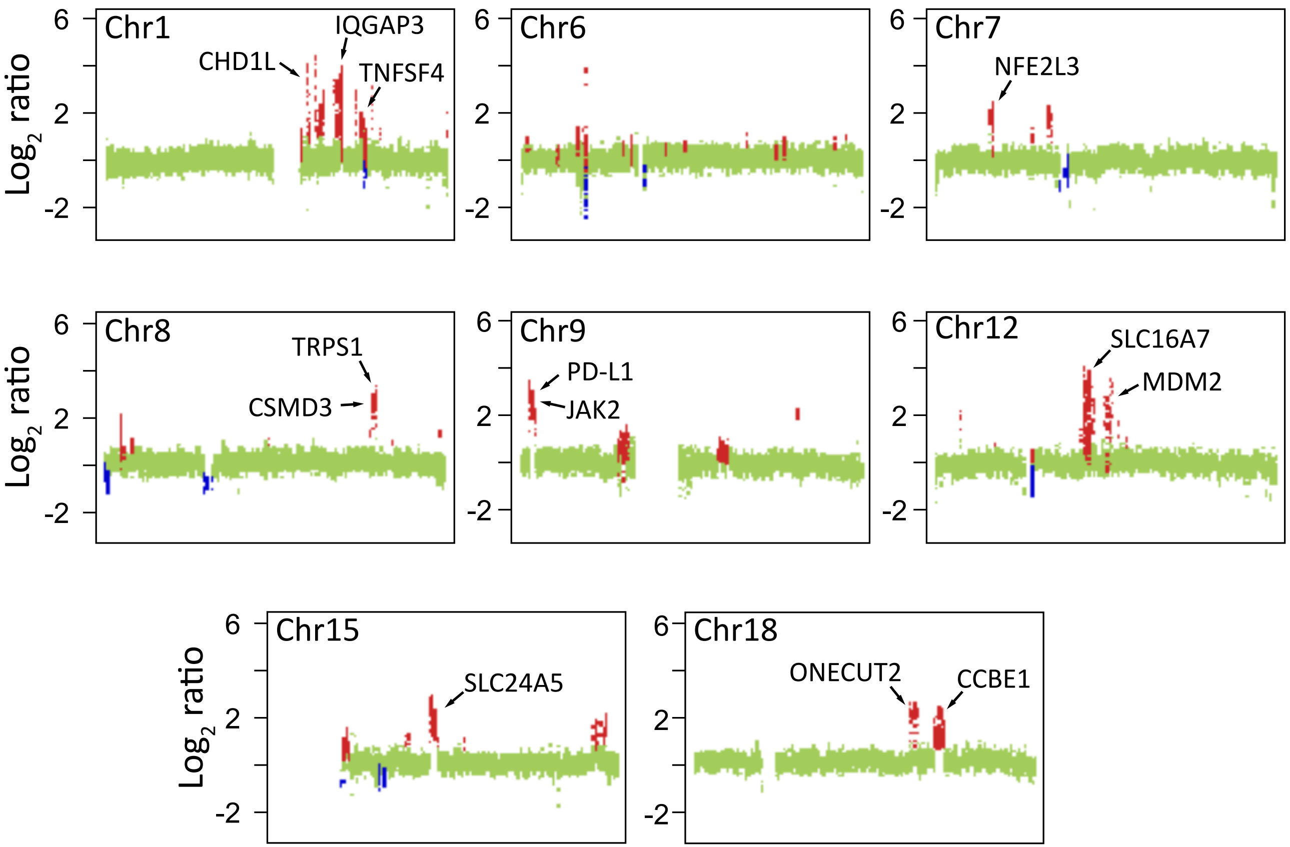


**Figure S7. CNV analysis of renal cancer tissue.** CNV analysis of chromosomes with abnormal amplification in renal cancer tissue. Gains and losses of copy number are shown in red and blue, respectively. Representative genes located in amplification region are marked with arrows.
