## Supplemental Fig 8 for "Robust identification of extrachromosomal DNA and genetic variants using multiple genetic abnormality sequencing (MGA-Seq)"

### Figure S8

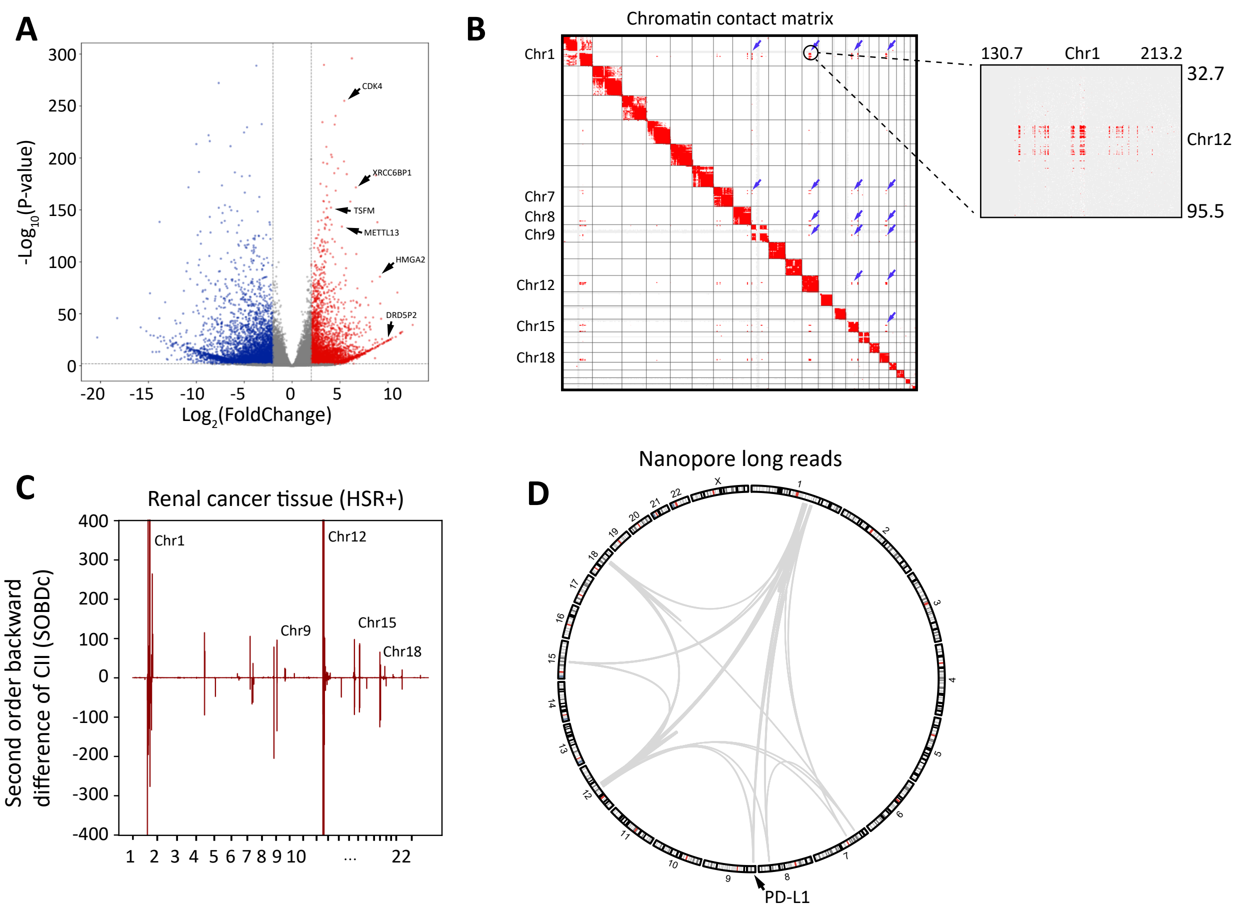


**Figure S8. Verification of inter and intra chromosomal interaction between focal amplification regions in renal cancer tissue by nanopore. (A)** Volcano plots of differential expression genes between renal cancer tissue and normal kidney tissue control. **(B)** Genome-wide chromatin contact matrix of renal cancer tissue. Potential HSR regions are marked with arrows in the figure. The inter-chromosomal contacts between the focal amplification regions and Chr1 and Chr12 are zoomed in. **(C)** The second order backward difference (SOBD) value across the genome of the renal cancer tissue in 100-kb bin size. **(D)** Validation of split reads and chromatin interactions across focal amplification regions with Nanopore long reads.
