## Supplemental Fig 9 for "Robust identification of extrachromosomal DNA and genetic variants using multiple genetic abnormality sequencing (MGA-Seq)"

### Figure S9

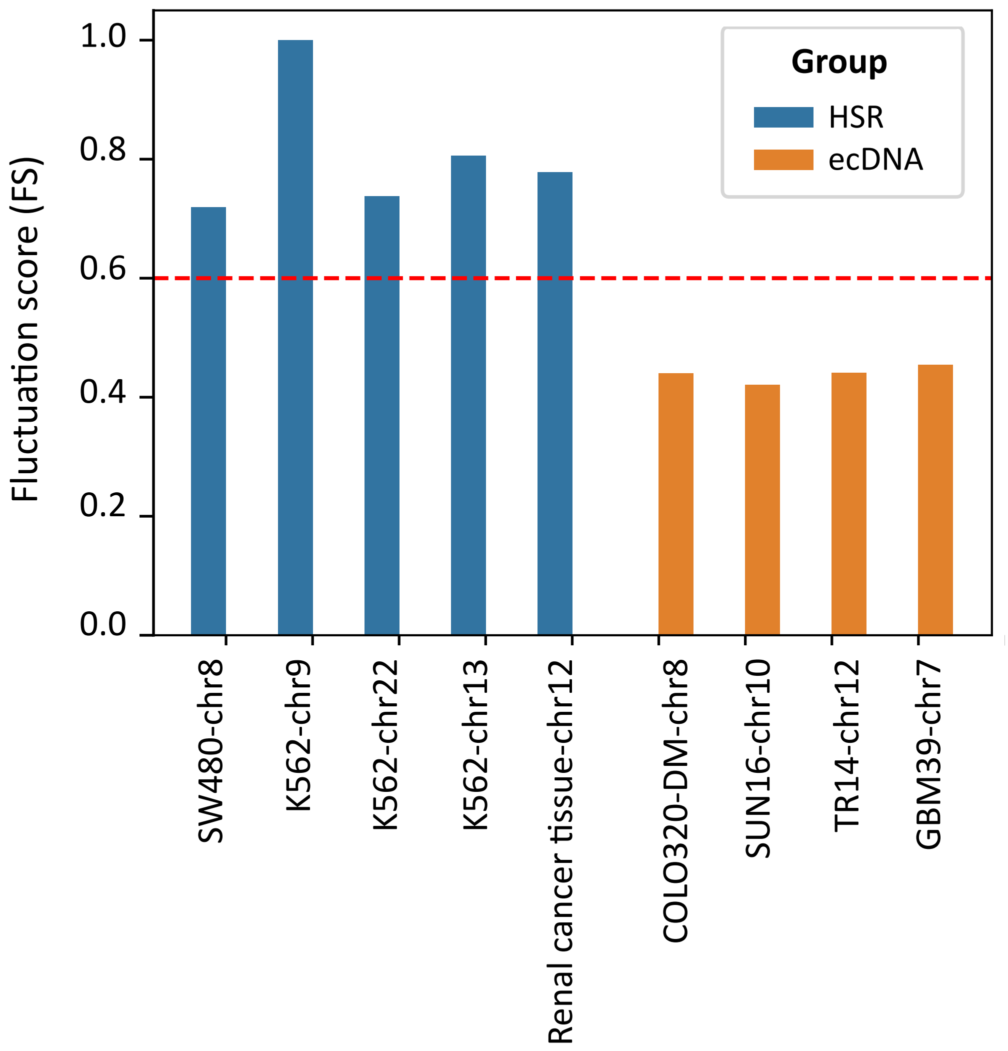


**Figure S9. Distribution of fluctuation score (FS) in different cell lines.** Blue bars indicate HSR-positive cell lines and yellow bars indicate ecDNA-positive cell lines. The Y axis represents the value of FS.
