## Supplemental Table 2 for "Robust identification of extrachromosomal DNA and genetic variants using multiple genetic abnormality sequencing (MGA-Seq)"

| CHR | START | END | SIZE(BP) | GENE |
|---|---|---|---|---|
| chr1 | 146,673,493 | 146,827,075 | 153,582 | CHD1L |
| chr1 | 147,137,879 | 147,203,090 | 65,211 | BCL6 |
| chr1 | 148,840,689 | 148,956,890 | 116,201 | DRD5P2 |
| chr1 | 149,794,421 | 149,843,524 | 49,103 | H4C14 |
| chr1 | 153,095,593 | 153,207,599 | 112,006 | SPRR2C |
| chr1 | 154,751,326 | 154,777,385 | 26,059 | KCNN3 |
| chr1 | 155,354,488 | 155,376,546 | 22,058 | ASH1L |
| chr1 | 156,497,604 | 156,514,786 | 17,182 | IQGAP3 |
| chr1 | 157,035,133 | 157,079,631 | 44,498 | ETV3L |
| chr1 | 157,291,307 | 157,464,725 | 173,418 | FCRL5、FCRL4、FCRL3 |
| chr1 | 158,315,548 | 158,734,921 | 419,373 | SPTA1 |
| chr1 | 167,929,100 | 167,955,539 | 26,439 | DCAF6 |
| chr1 | 168,408,796 | 168,922,040 | 513,244 | XCL1, XCL2, DPT, LINC00626 |
| chr1 | 171242769 | 173295748 | 2,052,979 | PRRC2C, MYOC, EEF1AKNMT, DNM3, FASLG, TNFSF18, TNFSF4 |
| chr1 | 183238351 | 183321805 | 83,454 | NMNAT2 |
| chr6 | 157731013 | 157735728 | 4,715 | TMEM242 |
| chr6 | 161025985 | 161073729 | 47,744 | LPA |
| chr7 | 25860032 | 26515412 | 655,380 | NFE2L3, CBX3, SNX10 |
| chr7 | 53290121 | 53708881 | 418,760 | |
| chr8 | 115412039 | 116285413 | 873,374 | CSMD3, TRPS1 |
| chr9 | 3908476 | 6438619 | 2,530,143 | JAK2, PD-L1, RCL1 |
| chr12 | 57,959,398 | 58,818,140 | 858,742 | KIF5A, DTX3, OS9, TSPAN31, TSFM, CTDSP2, GIHCG |
| chr12 | 59,353,303 | 59,563,937 | 210,634 | LRIG3 |
| chr12 | 59,980,377 | 60,217,376 | 236,999 | SLC16A7 |
| chr12 | 60,720,157 | 60,846,356 | 126,199 | |
| chr12 | 61,415,399 | 61,419,610 | 4,211 | |
| chr12 | 61,757,440 | 61,769,985 | 12,545 | |
| chr12 | 62,150,507 | 62,460,861 | 310,354 | TAFA2 |
| chr12 | 66,205,626 | 66,243,779 | 38,153 | HMGA2 |
| chr12 | 67,280,965 | 67,314,764 | 33,799 | RPSAP52 |
| chr12 | 67,907,648 | 67,949,103 | 41,455 | LINC02408 |
| chr12 | 68,217,256 | 68,315,666 | 98,410 | |
| chr12 | 68,399,408 | 68,419,703 | 20,295 | IFNG-AS1 |
| chr12 | 69,141,316 | 69,259,287 | 117,971 | SLC35E3, MDM2 |
| chr12 | 71,749,239 | 71,778,230 | 28,991 | LGR5 |
| chr15 | 45,899,960 | 48,282,442 | 2,382,482 | SQOC, SEMA6D, SLC24A5 |
| chr18 | 49,300,486 | 49,683,901 | 383,415 | |
| chr18 | 50,295,622 | 50,895,144 | 599,522 | DCC, MIR4528 |
| chr18 | 54,670,204 | 56,219,549 | 1,549,345 | ST8S1A3, ONECUT2, FECH, ATP8B1, NEDD4L, ALPK2 |
| chr18 | 56,508,853 | 57,185,058 | 676,205 | ZNF532, OACYLP, SEC11C, GRP, CPLX4, CCBE1 |

**Table S2. The regions of focal amplification in the renal cancer tissue.**
